# Extended floral longevity does not increase seed production under experimental pollinator decline

**DOI:** 10.64898/2026.07.13.738293

**Authors:** Gurleen K. Chana, Hanna Hickey, Christina M. Caruso

**Affiliations:** Department of Integrative Biology, University of Guelph, Guelph, Ontario, Canada

**Keywords:** floral longevity, floral traits, *Lobelia siphilitica*, natural selection, phenotypic selection, pollinator declines

## Abstract

**Premise of research:** In plant species that depend on pollinators to produce seeds, declines in pollinator populations should strengthen natural selection for floral traits that increase outcross pollen receipt by increasing the probability of pollinator visitation. One such trait is floral longevity, the amount of time that a flower is open and functional. If flowers that are open longer have a higher probability of pollinator visitation and this pollination benefit outweighs the resource cost of maintaining floral tissue, then pollinator decline should strengthen selection for extended floral longevity.

**Methodology:** To test whether pollinator decline could strengthen selection for extended longevity, we studied the pollinator-dependent species *Lobelia siphilitica*. We exposed *L. siphilitica* plants to either an ambient open-pollination treatment or a reduced open-pollination treatment that simulates pollinator decline. Within each pollination treatment, we measured floral longevity and seeds per fruit and used these data to estimate directional selection on longevity. This experiment was repeated twice.

**Pivotal results:** In one experiment, there was significant selection within both ambient and reduced pollination treatments for shortened (rather than extended) floral longevity. In the other experiment, selection on floral longevity within both ambient and reduced pollination treatments was not significantly different from zero.

**Conclusions:** We found no evidence of selection for extended floral longevity, regardless of the pollination environment. These results suggest that pollinator decline will not strengthen selection for extended longevity and thus that pollinator-dependent species such as *L. siphilitica* may not respond to pollinator decline by evolving extended floral longevity.

## Background

Populations of many pollinators are declining (Cameron et al. 2011; Cornelisse et al. 2025). As pollinators decline, so does the reliability of pollination services, such that plants receive less pollen and produce fewer seeds (Rodger et al. 2021; Artamendi et al. 2025). Pollinator decline should reduce the reliability of pollination services by affecting pollinator visitation rate rather than the fit between flower and pollinator. Consequently, pollinator decline should result in natural selection for floral traits that increase outcross pollen receipt by increasing the probability of pollinator visitation (Opedal 2021). Such selection should be particularly strong on floral traits of plant species that cannot autonomously self-pollinate and thus require pollinator visitation to produce seeds (i.e., pollinator-dependent species; Rodger et al. 2021). In these pollinator-dependent species, pollinator decline can reduce the mean absolute seed production in a population and select for floral traits that cause individual plants to produce more seeds relative to other plants in the population (Sletvold et al. 2016; Hossack and Caruso 2023; Brown and Caruso 2023). If the floral traits that are selected for are heritable, then they could evolve in response to pollinator decline (reviewed in Thomann et al. 2013).

One heritable (Spigler and Woodard 2019) floral trait of pollinator-dependent plant species that could be selected for as pollinators decline is extended floral longevity. Floral longevity is the amount of time that an individual flower is open and functional (Primack 1985). Extended floral longevity should increase outcross pollen receipt because flowers that are open longer have a higher probability of pollinator visitation, even if they are not more attractive or rewarding to pollinators (Rathcke 2003). If plants with flowers that are open longer receive more outcross pollen, then they should produce more seeds. However, whether plants with flowers that are open longer will produce more seeds depends not only on the pollination benefits of extended longevity, but also on the resource costs (Ashman and Schoen 1994; Schoen and Ashman 1995). Specifically, keeping flowers open longer can be costly if more carbon is allocated to support respiration by floral tissue (Williams et al. 1985; Galen et al. 1993) or if less carbon is fixed because unregulated water loss from floral tissue causes stomatal inhibition of photosynthesis (Lambrecht 2013; Roddy et al. 2024). If these costs of extended floral longevity reduce the carbon available to mature fertilized ovules, then flowers that are open longer could receive more outcross pollen yet produce fewer seeds. Consequently, whether extended floral longevity increases seed production and thus is selected for will depend on whether the pollination benefits of keeping flowers open longer outweigh the resource costs (Schoen and Ashman 1995).

Selection for extended floral longevity should be strengthened as pollinators decline because the pollination benefits of keeping flowers open longer are increased, but the resource costs are unchanged. Pollinator decline should increase the benefits of extended longevity because the amount of pollen deposited on a flower per day (i.e. the female fitness accrual rate; Ashman and Schoen 1994) is reduced when pollinators are scarce (e.g. Blair and Wolfe 2007), while the degree to which reproduction is pollen limited is increased (e.g. Sletvold and Agren 2016; Brown and Caruso 2023). In contrast, pollinator decline should not affect the costs of extended longevity because the availability of resources to mature fertilized ovules is determined not by the pollination environment but by the abiotic environment (e.g. Dudley et al. 2018). But despite the potential for pollinator decline to select for extended floral longevity, selection on longevity has yet to be estimated in any pollination environment.

To test whether pollinator decline could strengthen selection for extended floral longevity, we studied *Lobelia siphilitica* (Campanulaceae). *Lobelia siphilitica* is a relevant species for studying responses to pollinator decline because its primary pollinator (Bombus spp.; Macior 1967; Flanagan et al. 2011) is declining in portions of its range (Colla and Packer 2008; Cameron et al. 2011). *Lobelia siphilitica* should be vulnerable to a decline in pollination services because it is dependent on pollinators to produce seeds: female *L. siphilitica* flowers do not produce pollen, whereas hermaphroditic *L. siphilitica* flowers cannot enter the female phase until all pollen has been removed (i.e., complete dichogamy; Dudle 1999), eliminating the opportunity for autonomous self pollination (Beaudoin Yetter 1989). As expected given that it is a pollinator-dependent species, *L. siphilitica* that were exposed to experimentally simulated pollinator decline were pollen limited (Brown and Caruso 2023) and produced 18-21% fewer seeds than plants exposed to ambient pollination conditions (Brown and Caruso 2023; Hossack and Caruso 2023). Although open-pollinated *L. siphilitica* flowers can remain open for as long as 16 days (Lee and Caruso 2022b). this extended longevity incurs a resource cost: flowers hand pollinated on day 5 of the female phase produce 24-34% fewer seeds per fruit than flowers hand pollinated on day 1 of the female phase despite receiving the same pollen load, as expected if there is a cost of extended longevity (McCabe and Caruso 2026; Foster and Caruso 2022). But if open-pollinated *L. siphilitica* flowers with extended longevity receive more outcross pollen when pollinators are scarce, and this pollination benefit outweighs the resource cost, then pollinator decline should strengthen selection for extended floral longevity.

To determine the effect of pollinator decline on selection on floral longevity, we ran two experiments. In both experiments, we exposed *L. siphilitica* plants to either an ambient open-pollination treatment (where pollinator access to plants was unmanipulated) or a reduced open-pollination treatment (where pollinator access to plants was reduced by ∼50% to simulate pollinator decline; Lee and Caruso 2022b; Brown and Caruso 2023; Hossack and Caruso 2023). Within each treatment, we measured floral longevity and seeds per fruit to estimate directional selection on longevity. We compared these estimates of selection between pollination treatments within each experiment to answer the following question: Does pollinator decline strengthen selection for extended floral longevity?

## MATERIALS AND METHODS

### Study Species

*Lobelia siphilitica* is a self-compatible perennial wildflower that grows in wet meadows and woods throughout eastern North America (Johnston 1991 and references therein). *Lobelia siphilitica* populations flower from early August to late October (Miller and Stanton-Geddes 2007) and appear to have a unimodal flowering schedule (C. M. Caruso, personal observation). *Lobelia siphilitica* plants produce 3-cm-long blue to purple flowers (Caruso et al. 2003, 2010) that bloom acropetally along a racemose inflorescence. In *L. siphilitica*, the longevity of open-pollinated flowers ranges from 2-16 days (Lee and Caruso 2022b) and is not correlated with plant size estimated as final aboveground biomass (r ranges from −0.353 to 0.211 among treatment combinations, all P > 0.05; A. Whitehead and C. M. Caruso, unpublished data). Hermaphroditic *L. siphilitica* flowers are protandrous, with anthers and filaments fused into a tube that houses the style and stigma (Johnston 1991). Once *L. siphilitica* pollen is dispensed from the anthers by a pollinator, the style grows through the anther tube, at which point the stigma lobes reflex and become receptive (Ladd 1994). Although *L. siphilitica* plants generally produce a single inflorescence per flowering season, they can produce rosettes that overwinter and produce a raceme in the following year (Pigliucci and Schlichting 1995).

*Lobelia siphilitica* populations can contain both female and hermaphroditic individuals (i.e., gynodioecy; Beaudoin Yetter 1989). Although the frequency of female *L. siphilitica* within a population can vary from 0-100% (Caruso and Case 2007, 2013; Case and Caruso 2010; Appiah-Madson et al. 2022), the experimental population that we collected seeds from contained 31% females (Hossack and Caruso 2023).

### Germination and Growth

To generate plants for our experiment, we germinated seeds collected in 2020 from open-pollinated *L. siphilitica* plants in an experimental population (see Hossack and Caruso (2023) for more details). To induce germination, seed dormancy was broken using a dilute solution of chlorine bleach and ethanol (as in Dudle et al. 2001). On February 14-18 2022, seeds were planted into plug trays filled with ProMix BX (Premier Tech, Riviere-du-Loup, Quebec, Canada) and placed haphazardly on a greenhouse bench in the University of Guelph Phytotron (Guelph, Ontario, Canada). Six weeks later, seedlings (N = 200) were transplanted into 635 mL Deepots (Stuewe and Sons Inc., Tangent, Oregon, USA) filled with ProMix BX and randomly assigned a position on a greenhouse bench. Plants were watered using an automated drip irrigation system, fertilized with eight pellets of Nutricote 13-13-13 with micronutrients (Plant Products, Brampton, Ontario, Canada), and exposed to supplemental light (16-h days). *Lobelia siphilitica* plants (N = 87 females and 91 hermaphrodites) were transplanted into 8.7 L pots filled with ProMix BX prior to being moved into the field.

### Experimental Design and Data Collection

*Lobelia siphilitica* plants (N = 178) were moved into a fenced field site at the University of Guelph Arboretum (Guelph, Ontario, Canada; 43.542241, −80.220132). At the field site, plants were randomly assigned to a position in a 12 × 15 m grid. This grid contained four rectangular arrays of 45 plants (15 plants per row × 3 rows). Plants were watered as needed to maintain soil moisture at field capacity. Plants remained in the field from June 20 2022 until October 3 2022.

When the *L. siphilitica* plants at the field site began flowering, we used them for two pollination experiments. The two experiments were conducted using two sequentially produced samples of flowers on the same plants (e.g. Sahli and Conner 2011; Wassink and Caruso 2013). Experiment #1 was conducted from late June until mid-July, whereas Experiment #2 was conducted from late July until mid-August. For Experiment #1, we systematically assigned half of the plants to an ambient open-pollination treatment and half of the plants to a reduced open-pollination treatment that mimics pollinator decline. Plants in the ambient pollination treatment were unmanipulated. Plants in the reduced pollination treatment were covered with mesh pollinator exclusion bags for two days and then uncovered for two days, alternating for the duration of the experiment (i.e., reducing pollinator access by ∼50%). These treatments have been used in previous studies of the effect of pollinator decline on selection on floral traits of *L. siphilitica*, where they did not affect daily display size (Hossack and Caruso 2023; Brown and Caruso 2023) or flowering span (t_272_ = 1.505; P = 0.134; data from Hossack and Caruso (2022)). Pollination treatments were assigned separately for female and hermaphroditic plants, such that half of the plants of each sex were exposed to each treatment. Pollination treatments were switched for Experiment #2, such that plants assigned to the ambient pollination treatment in Experiment #1 were assigned to the reduced pollination treatment in Experiment #2 and vice versa. For Experiment #2 the sample size for was reduced to N = 148 because some plants had died or finished flowering.

To determine whether pollinator decline strengthens selection for extended floral longevity, we estimated the longevity and number of seeds per fruit for a sample of *L. siphilitica* flowers. On each plant in each experiment, we used colored craft wire to mark the pedicels of five fully expanded floral buds. We visited plants with marked buds daily to record the date of anthesis and the date of corolla wilt. We estimated floral longevity as the number of days from anthesis to corolla wilt (as in Lee and Caruso 2022b; McCabe and Caruso 2026). When marked flowers produced mature fruit, they were collected to estimate seeds per fruit. Because *L. siphilitica* seeds are small and numerous, we did not directly count the number of seeds per fruit. Instead, we established a relationship between seed mass and seed number by calculating the mean mass of 50 seeds from 75 randomly selected fruits. We then estimated the number of seeds per fruit by weighing them, dividing by the mean mass of 50 seeds, and multiplying by 50. Floral longevity and seeds per fruit were averaged across the five marked flowers on each plant in each experiment prior to analysis.

Our study differs from many other phenotypic selection studies (e.g. Sletvold and Agren 2016; Brown and Caruso 2023) in not measuring a suite of floral traits, Instead, we focused on measuring a single trait: floral longevity. We made this decision for two reasons. First, we measured only floral longevity because we were focused on testing a specific hypothesis as to why extended longevity should be selected for as pollinators decline. Second, we measured only floral longevity because it requires visiting flowers daily for as long as 2 weeks and thus is more time-consuming to measure than many other floral traits.

Our study also differs from many other phenotypic selection studies (reviewed in Caruso et al. 2019) in estimating selection on floral traits not via seeds per plant, but seeds per fruit. We estimated selection on floral longevity via seeds per fruit because we hypothesized that it would be the fitness component through which the resource costs and pollination benefits of extended floral longevity would be expressed. This hypothesis is supported by our previous work in *L. siphilitica* demonstrating that the resource cost of extended floral longevity reduces the number of seeds per fruit (Foster and Caruso 2022; McCabe and Caruso 2026). In addition, seeds per fruit and fruits per plant are not correlated in *L. siphilitica* (r ranges from 0.024-0.189, all P > 0.05, data from Hossack and Caruso (2022)), suggesting that there is not a trade-off between these two fitness components.

### Statistical Analyses

To determine whether there is selection for extended floral longevity, we estimated standardized directional selection differentials (S). Selection differentials were estimated using pooled data from female and hermaphroditic plants, but including sex as an independent variable in the model did not qualitatively affect our inferences about selection on floral longevity (table A1, available online). Directional selection differentials were estimated using regression. Separate regression models were used to analyze data from each experiment. The models included seeds per fruit as the dependent variable and floral longevity as the independent variable. Seeds per fruit was relativized by dividing by the mean seeds per fruit and floral longevity was standardized to a mean of 0 and variance of 1. Seeds per fruit was relativized and longevity was standardized separately for each pollination treatment (ambient vs. reduced), and separate regression models were used to estimate selection differentials in each treatment. In these models, the slope of the regression line estimates the standardized directional selection differential (Conner 1988), which represents both direct selection on longevity and indirect selection via correlated traits. If the floral longevity term was significant and the associated slope was positive, then we can conclude that there is selection for extended floral longevity.

To determine whether pollinator decline strengthens selection for extended floral longevity, we used analysis of covariance (ANCOVA). Separate ANCOVA models were used to analyze data from the two experiments. The models included relative seeds per fruit as the dependent variable; standardized floral longevity as the continuous independent variable; pollination treatment (ambient vs. reduced) as the categorical independent variable; and the floral longevity × pollination treatment term. If floral longevity × pollination treatment was significant and selection was positive (i.e., *S* > 0) and stronger in the reduced pollination treatment, then we can conclude that pollinator decline strengthens selection for extended floral longevity. In reaching this conclusion we assume that any difference in selection between treatments was caused by pollinators rather than by interactions with other biotic and abiotic factors.

To test whether (as in previous studies; Hossack and Caruso 2023; Brown and Caruso 2023) reducing pollination to simulate pollinator decline reduces the number of seeds produced by *L. siphilitica*, we used analysis of variance (ANOVA). Separate ANOVA models were used to analyze data from the two experiments. The models included seeds per fruit as the dependent variable; pollination treatment (ambient vs. reduced) and sex (female vs. hermaphrodite) as independent variables; and a treatment × sex interaction. If the pollination treatment term was significant and seeds per fruit was lower in the reduced pollination treatment, then we can conclude that (as in previous studies) experimentally simulating pollinator decline reduced the number of seeds produced by *L. siphilitica*.

To test whether (as in previous studies; Lee and Caruso 2022b) reducing pollination to simulate pollinator decline can affect the longevity of *L. siphilitica* flowers, we used ANOVA. Separate ANOVA models were used to analyze data from the two experiments. The models included floral longevity as the dependent variable; pollination treatment (ambient vs. reduced) and sex (female vs. hermaphrodite) as independent variables; and a treatment × sex interaction. If the pollination treatment term was significant, then we can conclude that (as in previous studies) experimentally simulating pollinator decline affected the longevity of *L. siphilitica* flowers.

Distributional assumptions for all models were tested by examining diagnostic plots of simulated residuals (DHARMa; 10.32614/CRAN.package.DHARMa). The assumptions for ANOVA models used to compare seeds per fruit (table 1) and floral longevity (table 2) between pollination treatments were generally met. In contrast, the assumptions for the regression models used to estimate selection differentials were often not met. To confirm that the results from these models were robust, we used bootstrapping to estimate standard errors for all selection differentials. The bootstrapped and parametric standard errors were very similar (results not shown) and as such we report and interpret the parametric estimates.

**Table 1.** Effect of experimental pollination reduction on the number of seeds per fruit produced by flowers on female and hermaphroditic *Lobelia siphilitica* plants.

| Source | Experiment #1 |  | Experiment #2 |  |
| --- | --- | --- | --- | --- |
| | $F_{1,174}$ | $P$ | $F_{1,144}$ | $P$ |
| Pollination | 0.388 | 0.534 | 11.076 | 0.001 |
| Sex | 3.093 | 0.080 | 0.010 | 0.920 |
| Pollination × Sex | 0.812 | 0.369 | 0.027 | 0.869 |

**Table 2.**
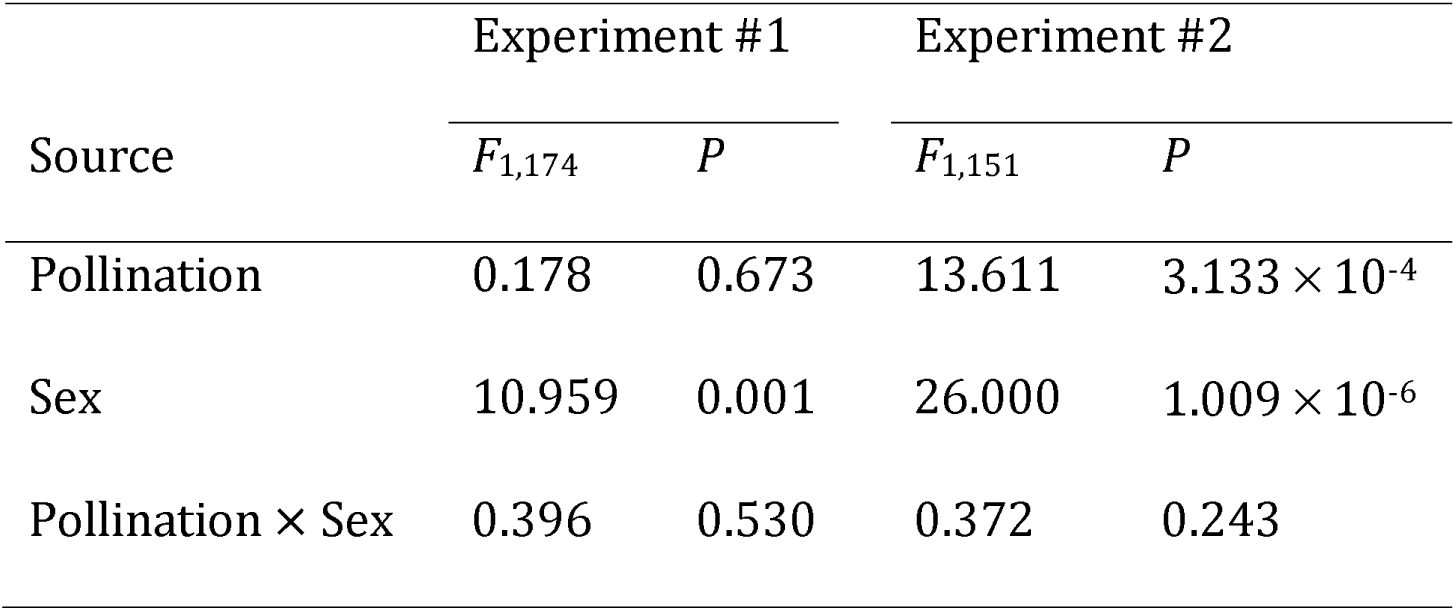
Effect of experimental pollination reduction on the longevity of flowers on female and hermaphroditic *Lobelia siphilitica* plants.

In addition to distributional assumptions, we confirmed the assumption that nonlinear selection on floral longevity was not significant (P > 0.05) by estimating standardized quadratic selection differentials (C) within each treatment and experiment (Table A2, available online). All statistical analyses were performed in R version 4.5.1 (R Core Team 2025).

## RESULTS

To determine whether pollinator decline strengthens selection for extended floral longevity, we compared selection differentials between ambient and reduced pollination treatments within each experiment. In Experiment #1, selection on longevity did not significantly differ between pollination treatments (non-significant floral longevity × pollination treatment term; *F*_1,174_ = 0.356, *P* = 0.551). Instead, in Experiment #1 there was significant selection for shortened (rather than extended) floral longevity in both the ambient (S_ambient_ ± 1 SE = −0.274 ± 0.096, *P* = 0.006; fig 1A) and reduced (*S_reduced_* ± 1 SE = −0.353 ± 0.089, *P* < 0.001; fig 1A) pollination treatments. In Experiment #2, selection on longevity also did not significantly differ between pollination treatments (non-significant floral longevity × pollination treatment term; *F*_1,144_= 0.005; P = 0.941). But in Experiment #2 (unlike in Experiment #1) selection on floral longevity was not significantly different from zero in either the ambient (*S_ambient_* ± 1 SE = −0.066 ± 0.072, *P* = 0.362; fig 1B) or reduced (*S_reduced_* ± 1 SE = −0.074 ± 0.088, P = 0.357; fig 1B) pollination treatment. This lack of selection for extended longevity across pollination treatments and experiments suggests that pollinator decline does not strengthen selection for extended floral longevity.

**Fig. 1.**
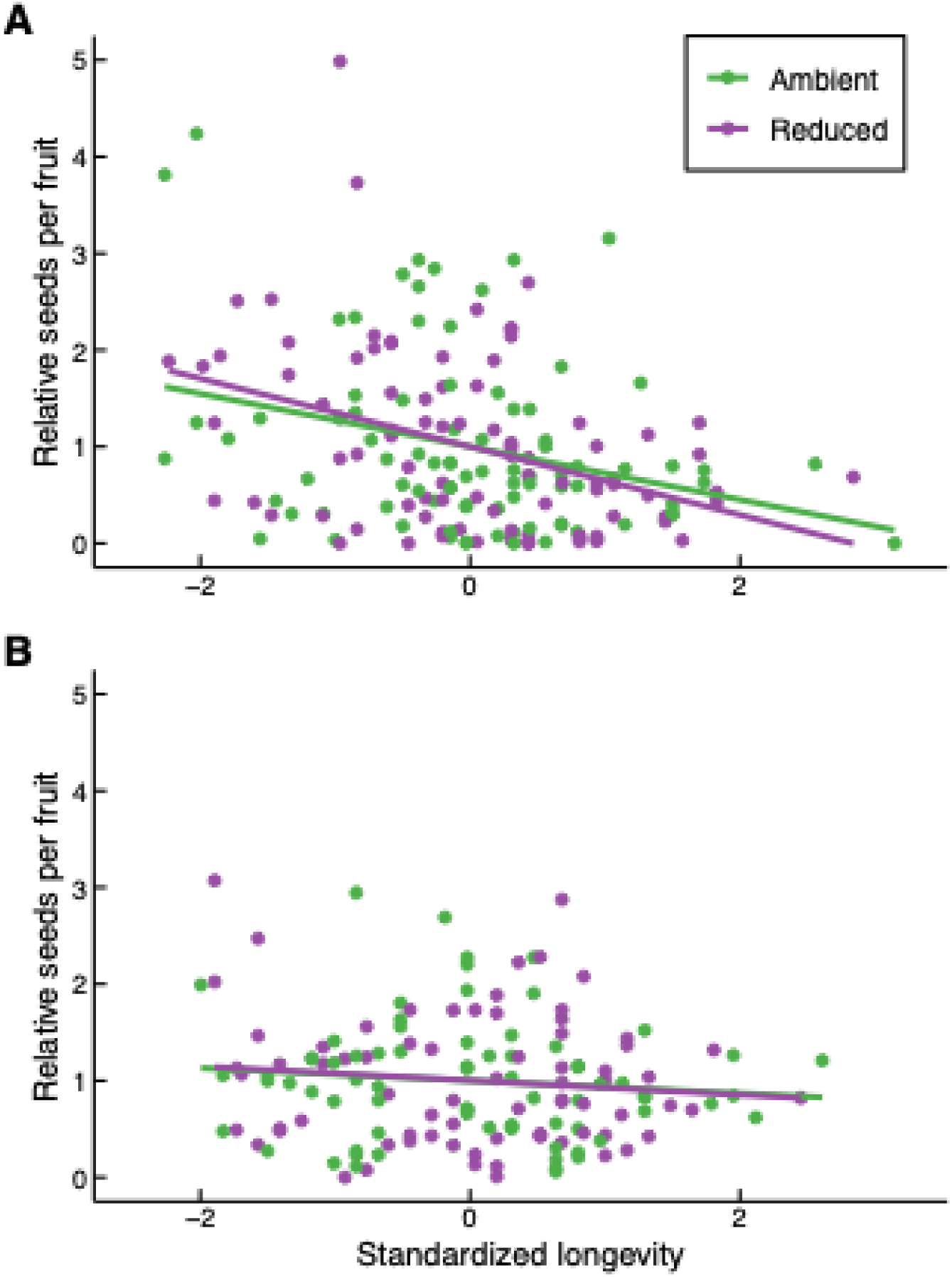
Relationship between floral longevity and seeds per fruit for *Lobelia siphilitica* plants exposed to ambient and reduced open-pollination treatments. (A) Experiment #1. (B) Experiment #2. Floral longevity was variance standardized and seeds per fruit was relativized within each pollination treatment.

To test whether reducing pollination to simulate pollinator decline reduces the number of seeds produced by *L. siphilitica*, we compared the number of seeds per fruit between pollination treatments. In Experiment #1, the number of seeds per fruit did not differ between ambient and reduced pollination treatments (non-significant pollination term; table 1; fig 2A). In contrast, within Experiment #2 the number of seeds per fruit was 30% lower in the reduced pollination treatment relative to the ambient treatment (significant pollination term; table 1; fig 2B). These results indicate that reducing pollination to simulate pollinator decline can (but does not always) reduce seed production in *L. siphilitica*.

**Fig. 2.**
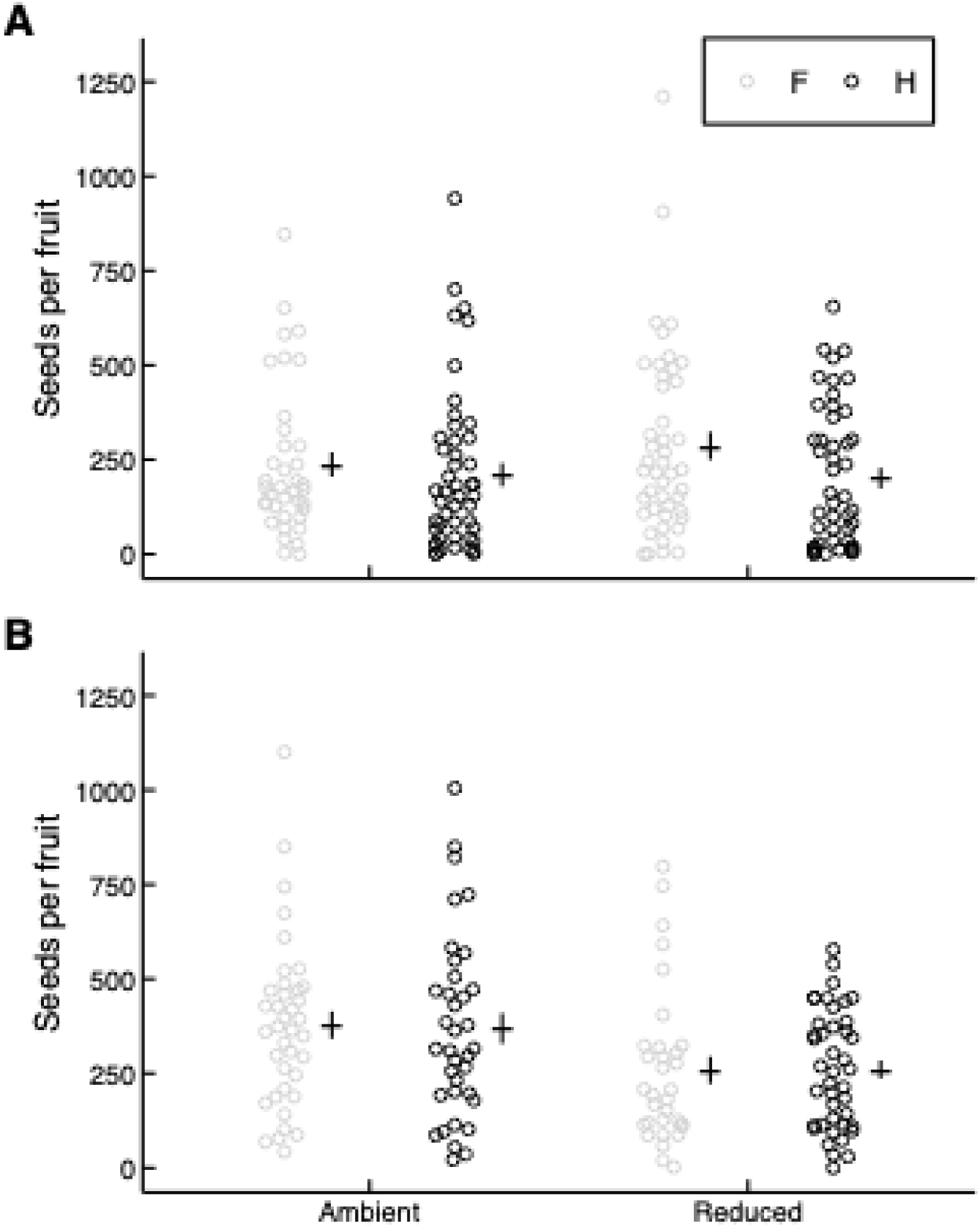
The number of seeds per fruit produced by female (F) and hermaphroditic (H) *Lobelia siphilitica* flowers exposed to ambient and reduced open-pollination treatments. The symbol to the right of the data points denotes the mean ± 1 SE for each combination of sex and pollination treatment. (A) Experiment #1. (B) Experiment #2.

To test whether reducing pollination to simulate pollinator decline can affect the longevity of *L. siphilitica* flowers, we compared floral longevity between pollination treatments. In Experiment #1, floral longevity did not significantly differ between ambient and reduced pollination treatments (non-significant pollination term; table 2; fig 3A). In contrast, in Experiment #2 longevity was 15.5% greater in the reduced pollination treatment relative to the ambient treatment (significant pollination term; table 2; fig 3B). These results indicate that reducing pollination to simulate pollinator decline can (but does not always) affect the longevity of *L. siphilitica* flowers.

**Fig. 3.**
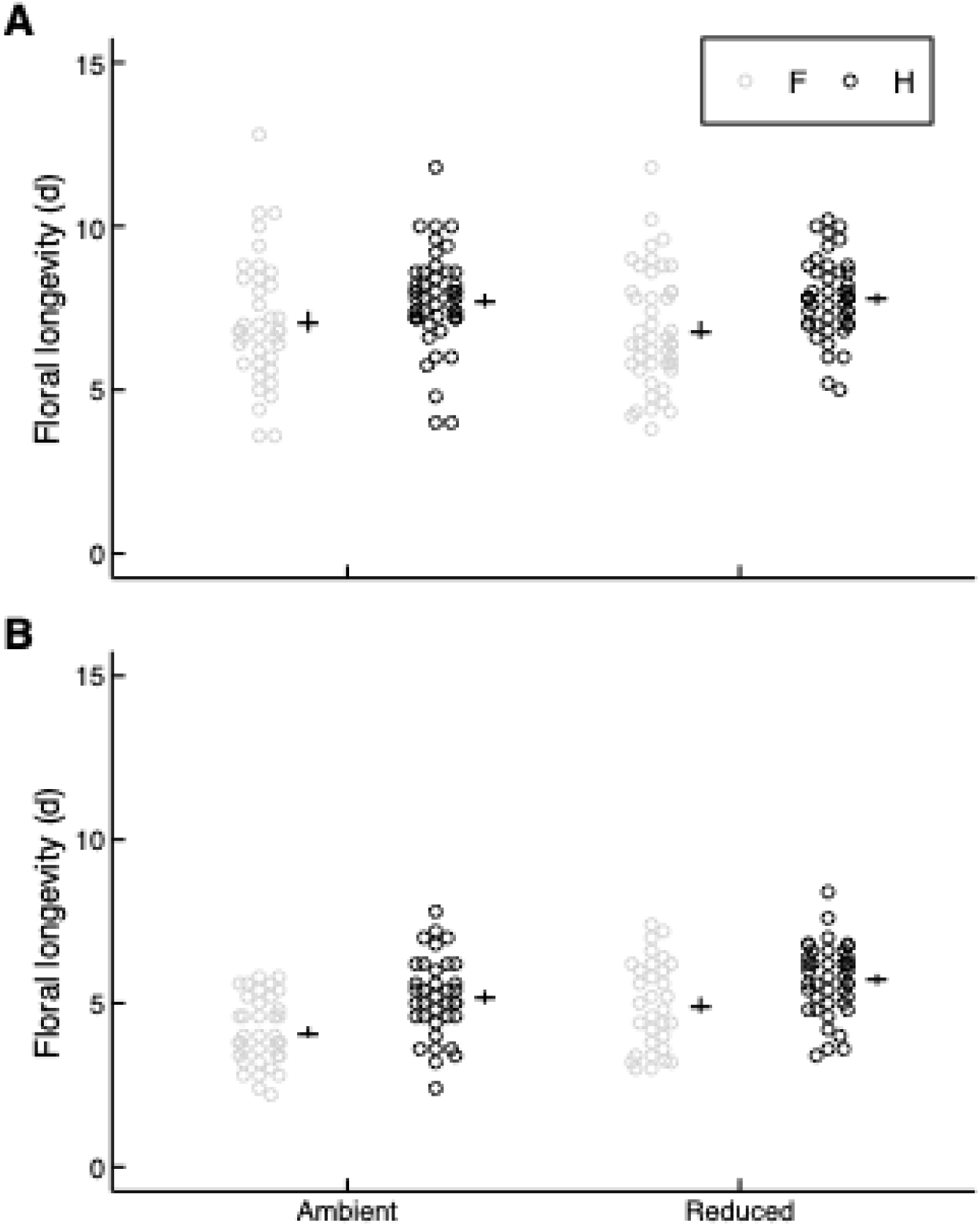
Longevity (in days) of female (F) and hermaphroditic (H) *Lobelia siphilitica* flowers exposed to ambient and reduced open-pollination treatments. The symbol to the right of the data points denotes the mean ± 1 SE for each combination of sex and pollination treatment. (A) Experiment #1 (CV_ambient_ = 22.83%; CV_reduced_ = 21.53%). (B) Experiment #2 (CV_ambient_ = 26.28%; CV_reduced_ = 23.22%).

## DISCUSSION

To determine whether pollinator decline strengthens selection for extended floral longevity, we did two experiments in which we simulated pollinator decline by reducing pollinator access to *L. siphilitica* plants. Regardless of the pollination treatment and experiment, we did not find evidence of selection for extended floral longevity. Instead, there was either significant selection for shortened longevity (Experiment #1; fig 1A) or no significant directional selection on longevity (Experiment #2; fig 1B), and the strength of this selection was not affected by experimentally simulated pollinator decline. These results do not support the hypothesis that pollinator decline, by increasing the pollination benefits of keeping flowers open longer, strengthens selection for extended floral longevity.

Our study is the first to estimate natural selection on floral longevity. Consequently, we do not know whether our results in *L. siphilitica*, where we found significant selection for shortened (Experiment #1; fig 1A) but not extended floral longevity, are typical or atypical. However, our results in *L. siphilitica* are consistent with the prediction (Ashman and Schoen 1994; Schoen and Ashman 1995) that selection for extended floral longevity will be limited by the resource cost of keeping flowers open longer. This cost is substantial in *L. siphilitica*: flowers hand-pollinated on day 5 of the female phase produce 24-34% fewer seeds per fruit than those hand-pollinated on day 1 (McCabe and Caruso 2026; Foster and Caruso 2022), indicating that the per-day cost of maintaining floral tissue is a 6%-8.5% decrease in the number of seeds per fruit. The per-day cost can be multiplied by the number of days that a flower is open to estimate the total resource cost of maintaining floral tissue across a flower’s lifespan (Schoen and Ashman 1995). The effect of this total resource cost on selection on floral longevity in *L. siphilitica* can be inferred by comparing Experiments #1 and #2. Because the longevity of flowers in Experiment #1 was ∼48% greater than in Experiment #2 (mean = 7.4 vs. 5 d; fig 3), the mean total resource cost of maintaining floral tissue should also be ∼48% greater in Experiment #1. And as expected if a high resource cost of keeping flowers open longer limits selection for extended longevity, there was significant selection for shortened floral longevity in Experiment #1 (fig 1A) but not Experiment #2 (fig 1B). But while costs of extended floral longevity are common (Ashman and Schoen 1997; Arathi et al. 2002; Castro et al. 2008; Marques and Draper 2012; Hildesheim et al. 2019; but see Spigler 2017), their magnitude varies among species (Foster and Caruso 2022), suggesting that the strength and direction of selection on floral longevity may also vary.

Contrary to our prediction that pollinator decline would strengthen selection for extended floral longevity, experimentally reducing pollinator access to simulate pollinator decline had no effect on selection on longevity in either experiment (fig 1). These results are not consistent with models of the evolution of floral longevity, which predict that poor (Schoen and Ashman 1995) or unpredictable (Xu and Servedio 2021) pollination environments should select for extended longevity. They are also not consistent with previous studies of *L. siphilitica* (Hossack and Caruso 2023; Brown and Caruso 2023), which found that experimentally reducing pollinator access to simulate pollinator decline strengthens selection for taller inflorescences and more vibrant petals (Hossack and Caruso 2023; Brown and Caruso 2023), both traits that can increase outcross pollen receipt by increasing the probability of pollinator visitation (e.g., Sletvold et al. 2016; Zu and Schiestl 2017). Instead, our results suggest that pollinator decline does not increase the pollination benefits of extended floral longevity in *L. siphilitica*, and thus there must be mechanisms that limit these benefits. For example, there could be limits on the pollination benefits of extended longevity because nectar rewards decline across the floral lifespan: if older flowers contain less nectar (e.g. Southwick and Southwick 1983; Devlin and Stephenson 1985; Wesselingh and Arnold 2000), then they could receive shorter pollinator visits and less outcross pollen (Manetas and Petropoulou 2000; Brandenburg et al. 2012) than younger flowers. More studies of the mechanisms that can limit the pollination benefits of extended floral longevity are needed to understand the conditions under which pollinator decline is more vs. less likely to select for extended longevity.

One limitation of our study is that we did not measure traits that are correlated with floral longevity in other species. These traits include flower size (e.g. Spigler and Woodard 2019) and daily display size (e.g. Harder and Johnson 2005). As such, we cannot rule out the possibility that there is direct selection for extended floral longevity in *L. siphilitica*, but that it is opposed by indirect selection for shortened longevity via correlated traits.

However, the longevity of *L. siphilitica* flowers is not phenotypically correlated with either flower size (*r* = 0.092, *N* = 67, *P* = 0.595; data from Hossack and Caruso (2022), Lee and Caruso (2022a)) or mean daily display size (Lee and Caruso 2022b). This lack of correlations suggests that any selection on flower size or daily display size in *L. siphilitica* would not cause indirect selection on floral longevity.

In conclusion, we did not find support for the hypothesis that pollinator decline, by increasing the pollination benefits of keeping flowers open longer, strengthens selection for extended floral longevity. Instead, there was either selection for shortened longevity (fig 1A) or no significant selection on longevity (fig 1B) in *L. siphilitica*, regardless of the pollination treatment. These results suggest that pollinator-dependent species such as *L. siphilitica* may not respond to pollinator decline by evolving extended floral longevity. The prediction that extended longevity will evolve in response to pollinator decline is compelling because flowers that are open longer have a higher probability of pollinator visitation even if they are not more attractive or rewarding to pollinators (Rathcke 2003). But although unusually extended floral longevity has evolved in some species (e.g., 33 days in *Telipogon peruvianus* (Orchidaceae); Martel et al. 2016), across angiosperms the median longevity is ∼4 days and the mode is ∼1 day (Song et al. 2022; Stephens et al. 2024), suggesting that the evolution of extended longevity is often limited. However, more studies that experimentally simulate pollinator decline and measure selection on floral longevity are needed to determine the extent to which our results in *L. siphilitica* can be generalized to other species.

## ACKNOWLEDGEMENTS

We thank M. Mucci and L. Illman for help in the greenhouse; H. Brazeau for help in the greenhouse and field; and two anonymous reviewers for comments on an earlier version of this manuscript. This work was supported by a Discovery Grant (awarded to CMC) and an Undergraduate Student Research Award (awarded to HH) from the Natural Sciences and Engineering Research Council of Canada. Data and code will be archived on Borealis (The Canadian Dataverse Repository).

---

**Table A1.** Standardized directional selection differentials (*S ± 1 SE (N)*) via seeds per fruit on floral longevity of *Lobelia siphilitica* exposed to ambient and reduced open-pollination treatments. These differentials were from models that included sex (female vs. hermaphrodite) as a fixed categorical factor.

|  | Ambient pollination | Reduced pollination |
| --- | --- | --- |
| Experiment #1 | $-0.272 \pm 0.100^{**} (86)$ | $-0.329 \pm 0.094^{***} (92)$ |
| Experiment #2 | $-0.074 \pm 0.080 (74)$ | $-0.083 \pm 0.084 (74)$ |

Note: Superscripts indicate whether a selection differential is significantly different from zero. $^{**} P < 0.01$ , $^{***} P < 0.001$ .

**Table A2.** Standardized quadratic selection differentials (*C ± 1 SE (N)*) via seeds per fruit on floral longevity of *Lobelia siphilitica* exposed to ambient and reduced open-pollination.

|  | Ambient pollination | Reduced pollination |
| --- | --- | --- |
| Experiment #1 | $0.051 \pm 0.059$ (86) | $0.033 \pm 0.066$ (92) |
| Experiment #2 | $-0.016 \pm 0.058$ (74) | $0.076 \pm 0.070$ (74) |

